# Unraveling molecular basis for reduced neuraminidase inhibitors susceptibility in highly pathogenic avian influenza A (H5N1) viruses isolated from chickens in India

**DOI:** 10.1101/2023.12.15.571865

**Authors:** Naveen Kumar, Richa Sood, Chhedi Lal Gupta, Sandeep Bhatia, Manoj Kumar, S. Nagarajan, C. Tosh, H.V. Murugkar, Aniket Sanyal

## Abstract

The increasing resistance cases in influenza viruses to different classes of antiviral drugs, necessitates the in-depth analysis of molecular interactions governing reduced susceptibility to these drugs. This study explores the molecular basis of neuraminidase inhibitors resistance in avian H5N1 influenza viruses identified in our previous research. Using comprehensive modeling and docking tools, we investigated two isolates—A/chicken/India/85459/2008 (N294S) and A/chicken/WestBengal/142121/2008 (E119A + I117V). The N294S mutation conferred oseltamivir resistance while retaining zanamivir susceptibility, whereas the E119A + I117V mutations led to zanamivir resistance while reducing oseltamivir susceptibility. Molecular interactions analysis unveiled varied fitness levels, hydrogen bonding, and affinity transitions associated with N294S and E119A + I117V mutations. This study provides crucial insights into molecular interactions responsible for reduced susceptibility to neuraminidase inhibitors, which is essential for optimizing antiviral strategies and pandemic preparedness.

---

With the growing concerns on acquisitions of resistance in influenza viruses to currently available antiviral drugs, monitoring of susceptibility in influenza viruses becomes critical for risk assessment and pandemic preparedness. As of right now, the Food and Drug Administration (FDA) has approved the two classes of antiviral drugs: M2 ion channel blockers, such as adamantanes (amantadine and rimantadine), and neuraminidase (NA) inhibitors (NAIs; oseltamivir, zanamivir, and peramivir) for the treatment of influenza virus infections. Nevertheless, adamantanes are no longer used to treat human influenza infections due to widespread emergence of adamantane-resistant strains (Nelson et al., 2009). As a result, NAIs are the only antiviral drugs used to treat influenza virus infections. NAIs target the highly conserved residues in NA proteins’ active sites. These active sites constitute eight catalytic residues (R118, D151, R152, R224, E276, R292, R371, and Y406) that interact directly with the NAIs, and eleven framework residues (E119, R156, W178, S179, D/N198, I222, E227, H274, E277, N294, and E425) that stabilize the active-site structure. Several *in vivo* and *in vitro* studies have demonstrated that emergence of NAI resistance is associated with mutations in these actives sites (Boltz et al., 2010; Hurt et al., 2009,2010; Ilyushina et al., 2010; Le et al., 2005; Naughtin et al., 2011; Nguyen et al., 2013; Sood et al., 2018; Bialy and Shelton, 2020; Wang et al., 2023). The influenza virus’s resistance/reduced susceptibility to NAIs is frequently assessed using the fluorescence-based NA inhibition assay by determining the 50% inhibitory concentration (IC_50_) of the NAIs to the influenza viruses. By measuring the IC_50_ fold change of an individual virus isolate compared to its subtype/clade specific median IC_50_ values, influenza viruses can be classified as resistant or susceptible to NAIs. If the fold change is less than 10, it is considered susceptible to the NAI tested, while 10- to 100-fold change is considered reduced susceptibility, and more than 100-fold change is considered highly reduced susceptibility to NAIs.

We previously identified oseltamivir and zanamivir resistance with established markers of resistance (E119A and N294S) and markers of reduced susceptibility (I117V) in five avian highly pathogenic avian influenza (HPAI) H5N1 clade 2.2 viruses using the fluorescence-based NA inhibition assay (Sood et al., 2018). However, the molecular basis conferring reduced susceptibility of these viruses to NAIs has not been investigated.

In order to unravel the molecular interactions responsible for NAI resistance/reduced susceptibility, two H5N1 isolates, A/chicken/India/85459/2008 (Oseltamivir IC_50_ = 307.42, Zanamivir IC_50_ = 4.30, N294S) and A/chicken/WestBengal/142121/2008 (Zanamivir IC_50_ = 203.0, Oseltamivir IC_50_ = 65.0, E119A + I117V) were selected for molecular interaction investigation. The N294S mutation in the NA protein (Uniprot ID: M4SYW9) of A/chicken/India/85459/2008 conferred resistance to oseltamivir (∼126-fold change in IC_50_ as compared to median IC_50_ of clade 2.2 viruses), while it remained susceptible to zanamivir (∼3-fold change). E119A + I117V mutations in NA protein (Uniprot ID: E2E1W8) of A/chicken/WestBengal/142121/2008, on the other hand, demonstrated resistance to zanamivir (∼125-fold change in IC_50_ as compared to median IC_50_ of clade 2.2 viruses) and reduced susceptibility to oseltamivir (∼27-fold change). Using comprehensive molecular modelling and docking computational tools, we elucidated the molecular interactions responsible for this varied susceptibility of influenza viruses harboring different set of NA mutations to NAIs.

To achieve this, we first built three-dimensional (3D) models of NA proteins of these two viruses on a protein databank template (PDB ID: 4b7q) via homology modeling using MODELLER 9v4 program (Sali and Blundell, 1993; Gupta et al., 2014). The 3D models with lowest Discrete Optimized Potential Energy (DOPE) scores were selected and their quality was further checked by ProSA (z-score of the input structure should be within the range of scores typically found for native proteins of similar size) (Wiederstein and Sippl, 2007) and ProQ (correct models should have LG score > 1.5 and MaxSub score > 0.1) servers (Wallner and Elofsson, 2003). The overall quality of 3D models of these proteins generated in this study were good as estimated by ProSA (Z scores were close to experimentally determined protein structure) and ProQ servers (LG scores > 2.5 and MaxSub scores > 0.3) (Supplementary Table 1).

In the docking experiment, ligands, Oseltamivir (PubChem CID: 78000) and Zanamivir (PubChem CID: 60855) were first corrected with Auto Edit Ligand and subsequently docked on the 3D models of the NA proteins, M4SYW9 (A/chicken/India/85459/2008) and E2E1W8 (A/chicken/West Bengal/142121/2008) using the GOLD (Genetic Optimization for Ligand Docking) v5.2 (Wu et al., 2003). The GOLD created the optimal position for interacting molecules by employing a genetic algorithm, and the top three poses with root-mean-square deviations (RMSDs) less than 1.5 Aº were chosen. The optimal docked ligand pose was chosen on the basis of CHEMPL score. Using the Discovery Studio visualizer tool, the interactions of docked complexes were examined, and an estimate of the Gold Fitness Score (GF score) was made. The ligand’s fitness to the protein active site pocket is represented by the GF score, which represents protein-ligand hydrogen bond energy, protein-ligand van der Waals energy, ligand internal energy and ligand torsional strain energy.

Molecular docking analyses revealed that the oseltamivir-M4SYW9 complex (GF score = 42.62, 1H-Arg273) exhibited a lower fitness level and fewer interactions, particularly in terms of hydrogen bonding, compared to the oseltamivir-E2E1W8 complex (GF score = 47.26, 4H-Arg98, and Arg348) (Supplementary Table 2). This disparity in interaction quality may explain the higher IC_50_ values observed for M4SYW9 in contrast to E2E1W8. Similarly, the zanamivir-susceptible isolate, A/chicken/India/85459/2008 (M4SYW9) demonstrated superior fitness and a higher level of hydrogen bonding (GF score = 54.62, 10H-Arg273, Ser275, Tyr382) compared to the zanamivir-resistant isolate, A/chicken/WestBengal/142121/2008 (E2E1W8) (GF score = 52.11, 2H-unknown) (Figure 1).

**Figure 1.**
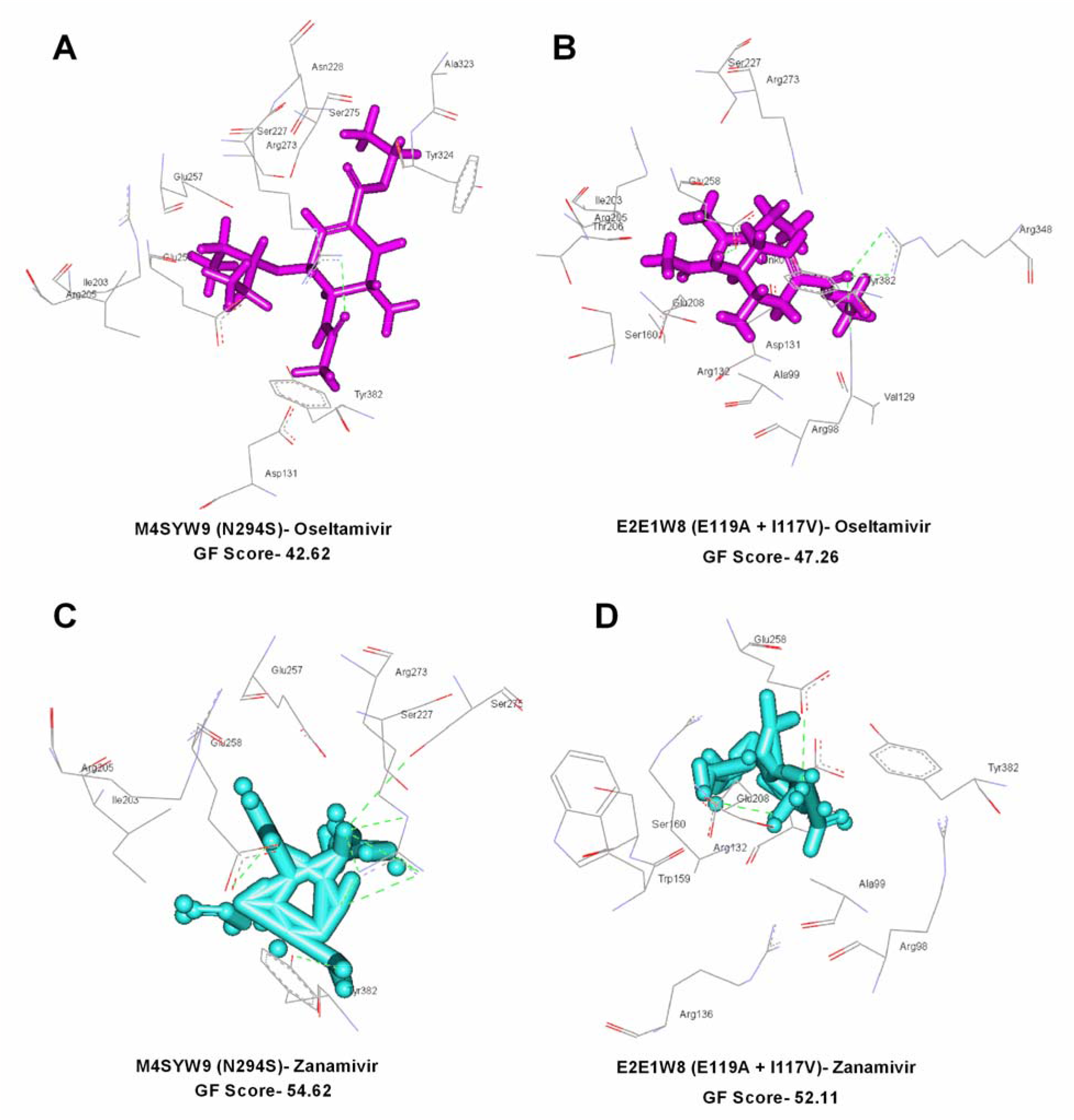
Molecular docking analyses of resistant and susceptible highly pathogenic avian influenza A (H5N1) viruses. The 3D models of these two isolates were docked with oseltamivir (A & B) and zanamivir (C & D) using GOLD v5.2 and visualized by discovery studio visualizer tool. Effect of NA substitutions within two H5N1 isolates, A/chicken/India/85459/2008 (N294S, Uniprot ID: M4SYW9) and A/chicken/WestBengal/142121/2008 (E119A + I117V, Uniprot ID: E2E1W8) on oseltamivir (A & B) and zanamivir (C & D) IC_50_ values was correlated with Gold Fitness Score and hydrogen bonding. Active site residues are displayed in stick form and hydrogen bonding is depicted in green dotted lines.

Furthermore, a comprehensive dissection of molecular interactions transitions due to NA-N294S and NA-I117V + E119A mutations was conducted. The substitution of asparagine at position 294 with serine minimally impacted the binding affinity of A/chicken/India/85459/2008 (M4SYW9) to zanamivir. However, the combined substitution of glutamic acid with aspartic acid and isoleucine with valine in A/chicken/WestBengal/142121/2008 (E2E1W8) led to the loss of hydrogen bonds with the amine group of Arg 273, the hydroxyl group of Ser 275, and the hydroxyl group of Tyr 382 (N1 numbering). This resulted in approximately a 47-fold reduction in binding affinity to zanamivir, as supported by a decreased GF score, compared to M4SYW9. Concerning oseltamivir, while both N294S and I117V + E119A mutations individually contributed significantly to the reduction of binding affinity, the N294S mutation was more potent in reducing hydrogen bond interactions with the NA active site residues. Specifically, oseltamivir could only form one hydrogen bond with the amine group of Arg273 in the presence of the N294S mutation (Supplementary Table 2). However, with the combined I117V + E119A mutation, oseltamivir formed four hydrogen bonds with Arg 98 and Arg 348, exhibiting a comparatively better GF score.

This molecular interaction study provides valuable insights into understanding the molecular basis of resistance to neuraminidase inhibitors (NAIs) influenced by various amino acid substitutions. Furthermore, it is evident that the impact of different mutations within the neuraminidase of influenza A virus varies widely depending on the subtype, clade, or the specific NAI (oseltamivir and zanamivir). Understanding these molecular intricacies is crucial for refining antiviral strategies and enhancing pandemic preparedness.

## Supporting information

Z scores were close to experimentally determined protein structure) and ProQ servers (LG scores > 2.5 and MaxSub scores > 0.3)

## Acknowledgments

The authors are thankful to the Indian Council of Agricultural Research (ICAR), New Delhi, India and the Director, ICAR-NIHSAD, Bhopal, India for providing infrastructural facilities to carry out this study. This study was funded in part by the Science and Engineering Research Board grant - CRG/2020/001274.

## Notes

### Competing Interest Statement

The authors have declared no competing interest.

