## Supplementary material for "Unraveling molecular basis for reduced neuraminidase inhibitors susceptibility in highly pathogenic avian influenza A (H5N1) viruses isolated from chickens in India": Z scores were close to experimentally determined protein structure) and ProQ servers (LG scores > 2.5 and MaxSub scores > 0.3)

**Supplementary Table 1:** Quality validation of three-dimensional (3D) models of NA proteins of H5N1 viruses

| **Uniprot ID** | **ProSA Server**  **(Z score)** | **ProQ server** | |
| --- | --- | --- | --- |
|  |  | **LG Score** | **MaxSub Score** |
| E2E1W8 (A/chicken/West Bengal/142121/2008) | -4.84 | 2.689 | 0.506 |
| M4SYW9 (A/chicken/India/85459/2008) | -4.90 | 3.003 | 0.318 |
| Template (4b7q) (experimentally determined structure) | -5.18 | 1.503 | 0.448 |

**Supplementary Table 2:** Comprehensive dissection of molecular interactions between neuraminidase and neuraminidase inhibitors using GOLD (Genetic Optimization for Ligand Docking) v5.2

| **Neuraminidase inhibitors** | **Uniprot ID** | **Gold Fitness Score** | **Residues making hydrophobic contact** | **No. of Hydrogen Bond formed** | **Hydrogen Bonding** | | |
| --- | --- | --- | --- | --- | --- | --- | --- |
|  |  |  |  |  | **Donor atoms** | **Acceptoratoms** | **Distance (Å)** |
| Oseltamivir | **M4SYW9** | 42.62 | Asp131  Ile203  Arg205  Ser227  Asn228  Glu257  Glu258  Arg273  Ser275  Ala323  Tyr324 | 1 | ARG273 :NH2 | UNK0 :O4 | 3.19 |
|  | **E2E1W8** | 47.26 | Arg98  Ala99  Val129  Asp131  Arg132  Ser160  Ile203  Arg205  Thr206  Glu208  Ser227  Glu258  Arg273  Arg348  Tyr382 | 4 | ARG98 :NH1 | UNK0 :O3 | 3.09 |
|  |  |  |  |  | ARG348 :NH1 | UNK0 :O3 | 2.96 |
|  |  |  |  |  | ARG348 :NH2 | UNK0 :O3 | 3.05 |
|  |  |  |  |  | UNK0 :H30 | GLU258: OE2 | 2.75 |
| Zanamivir | **M4SYW9** | 54.62 | Ile203  Arg205  Ser227  Glu257  Glu258  Arg273  Ser275  Tyr382 | 10 | ARG273 :NE | UNK0 :O2 | 3.02 |
|  |  |  |  |  | ARG273 :NH1 | UNK0 :O1 | 2.87 |
|  |  |  |  |  | ARG273 :NH1 | UNK0 :O2 | 2.68 |
|  |  |  |  |  | ARG273 :NH2 | UNK0 :O1 | 3.03 |
|  |  |  |  |  | ARG273 :NH2 | UNK0 :O2 | 2.87 |
|  |  |  |  |  | ARG273 :NH2 | UNK0 :O3 | 2.85 |
|  |  |  |  |  | SER275 :OG | UNK0 :O2 | 3.09 |
|  |  |  |  |  | TYR382 :OH | UNK0 :O6 | 2.52 |
|  |  |  |  |  | UNK0 :O5 | GLU258 :OE1 | 2.72 |
|  |  |  |  |  | UNK0 :O5 | GLU258 :OE2 | 2.83 |
|  | **E2E1W8** | 52.11 | Arg98  Ala99  Asp131  Arg132  Arg136  Trp159  Ser160  Glu208  Glu258  Tyr382 | 2 | UNK0 :N10 | GLU258 :OE2 | 2.86 |
|  |  |  |  |  | UNK0 N11 | GLU208 :OE2 | 3.18 |
